# Immuno-Proteomic Age of Cardiovascular Etiology (iPACE) Signature Predicts Cardiovascular Risk in Elderly Patients

**DOI:** 10.64898/2026.10.27.762320

**Authors:** Jordan J Baechle, Jennifer Arthur, Khiem Nguyen, Fei Wu, Matias Fuentealba, Kirk U. Knowlton, Stacey Knight, Benjamin Horne, Nazish Sayed, Holden Maecker, Daniel A. Winer, Nir Barzilai, Mark M. Davis, Francois Haddad, David Furman

## Abstract

Despite advances in clinical care, cardiovascular disease (CVD) remains the leading cause of disability and mortality worldwide. While age alone is a primary risk factor for CVD, accumulating evidence suggests the immune system influences the initiation and progression of CVD pathophysiology, particularly in heart failure with preserved ejection fraction (HFpEF) and atherosclerotic CVD (ASCVD). Here, using two independent cohorts of older adults, we develop a thirty-protein signature of age based on serum immune proteomics, which correlates with HFpEF and ASCVD risk and mortality (Immuno-Proteomic Age of Cardiovascular Etiology, iPACE). We find aging-related circulating immune proteins associated with ASCVD and HFpEF-specific resilience and susceptibility that may be used as potential prognostic, diagnostic, or therapeutic targets.

## Introduction

Despite advances in treatment and prevention, cardiovascular disease (CVD) remains the leading cause of death [1]. By 2030, approximately 20% of the population will be 65 years or older, and in this age group, CVD will result in 40% of all deaths [2]. Although there are many well-characterized modifiable risk factors for the development of CVD (including type 2 diabetes, obesity, tobacco use and high blood pressure and cholesterol), chronological age (cAge) is the most influential and unmodifiable [3].

Although the rate of chronological aging is solely time-dependent, biological aging is variable and dependent on the cumulative trajectory of aging hallmarks over time [3-5]. Among the CVDs, heart failure with preserved ejection fraction (HFpEF) exhibits profound interconnectedness to aging and chronic inflammation and remains poorly understood with no available treatment options [6]. Therefore, understanding the relationship between cAge, biological age, HFpEF, and other CVDs, including atherosclerotic CVD (ASCVD), is paramount in characterizing the integral etiology prerequisite to advancing treatment and prevention. Here we develop a biological age metric, referred to as iPACE (Immuno-Proteomic Age of Cardiovascular Etiology), using serum immune proteomics and demonstrate it as an effective predictor of HFpEF and ASCVD (Figure 1)

**Figure 1.**
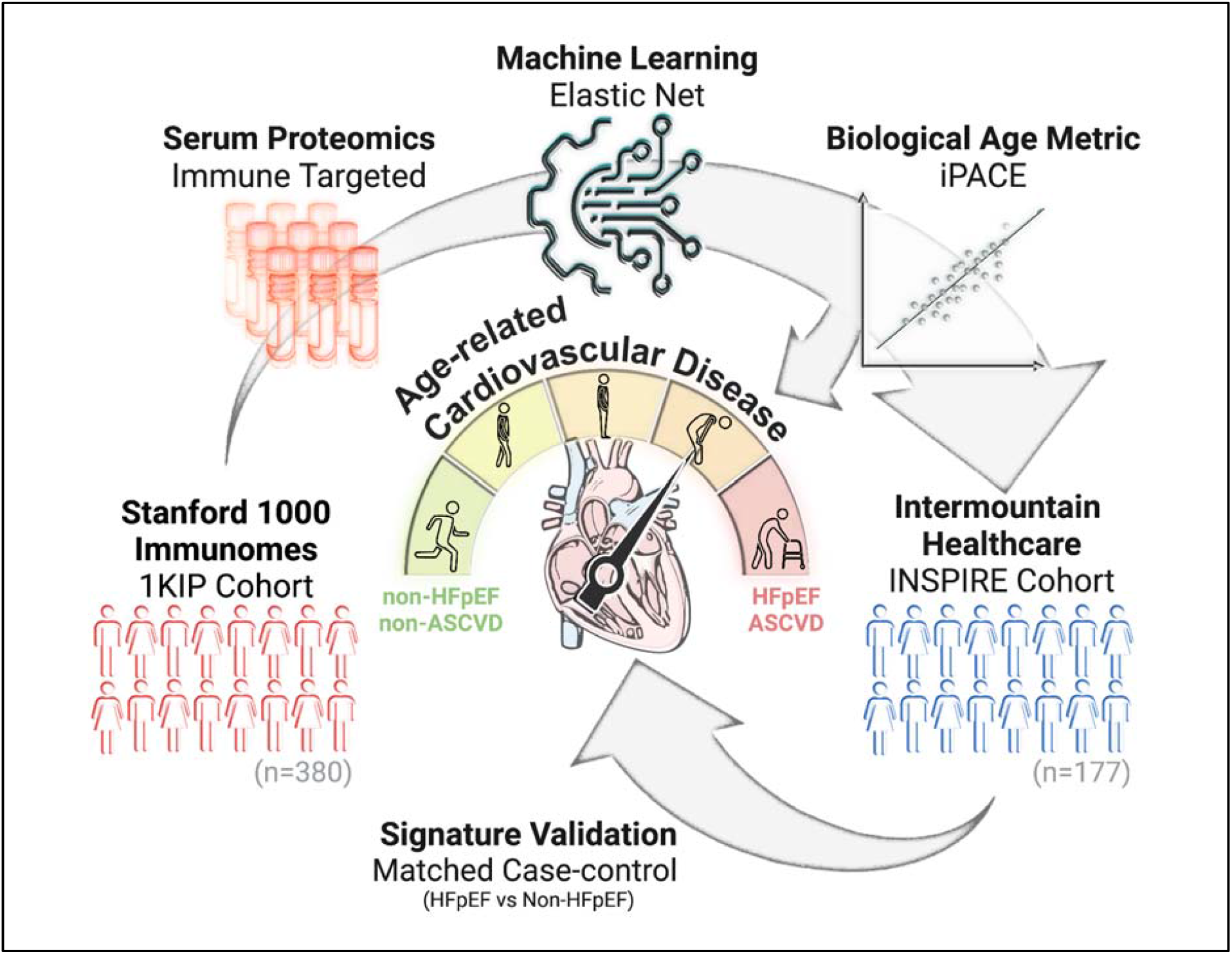
Study summary. Patients >60 years old from the Stanford 1000 Immunomes Project (1KIP) cohort. Targeted immune proteomics were used to generate a biological age metric using elastic net penalized regression. The biological age metric (iPACE = Immuno-Proteomic Age of Cardiovascular Etiology) was predictive of ASCVD (atherosclerotic cardiovascular disease) in 1KIP cohort. iPACE signature was validated using the INtermountain Healthcare Biological Samples Collection Project and Investigational Registry (INSPIRE) cohort as a predictor of HFpEF (heart failure with preserved ejection fraction) and ASCVD.

## Results

### Stanford 1000 Immunomes Project (1KIP) Cohort - Training Dataset

This study utilized the subset of the Stanford 1000 Immunomes Project (1KIP) cohort, which was followed annually over ten years using blood draws, medical records, and surveys to study age-related immune system changes. Only those patients 60 years or older were included in the current study. Further description of the cohort can be found at https://med.stanford.edu/1000immunomes.html.

### INSPIRE Cohort - Testing Dataset

The INSPIRE cohort is a subset of the INtermountain Healthcare Biological Samples Collection Project and Investigational REgistry for the Ongoing Study of Disease Origin, Progression, and Treatment study (INSPIRE). Patients 60 years or older diagnosed with HFpEF and case-matched controls were included in the study. Further description of the study can be found at https://clinicaltrials.gov/ct2/show/NCT02450006.

### Prediction of Chronological Age (cAge) and iPACE from Circulating Biomarkers

To identify predictors of cAge, cAge was regressed on circulating biomarkers of 1KIP participants (training dataset) using Elastic Net penalization. Data from the first and subsequent years of the study were used to train the model and evaluate its robustness, respectively. Leave-one-out cross-validation determined optimal tuning parameter ***λ*** and α from a 100x100 grid. The model identified thirty proteins predictive of cAge (Figure 2A). iPACE is the model’s predicted age. “iPACE residual” is the residual of iPACE and cAge adjusted for sex and used to quantify the degree of normal (iPACE residual=0), premature (iPACE residual>0), and delayed (iPACE residual<0) aging phenotypes.

**Figure 2.**
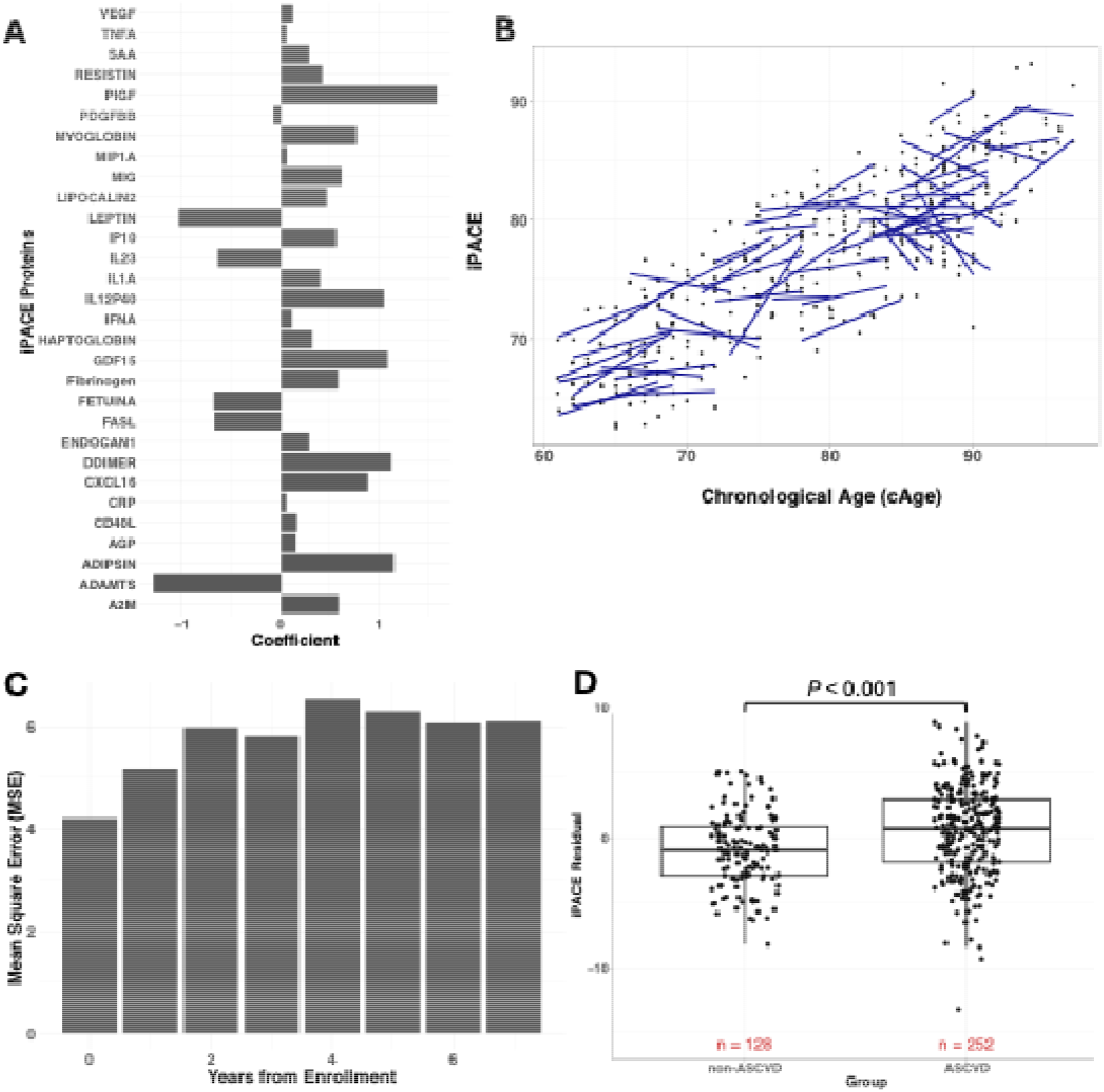
iPACE Model Discovery. Stanford 1000 Immunomes Project (1KIP) [training dataset] results: A) thirty-protein signature with weighted coefficients (iPACE) predictive of chronological age, B) Individual participant iPACE trajectories calculated by the slope of iPACE and cAge, C) iPACE mean square error according to year from enrollment, D) iPACE residual compared between patients with a diagnosis of atherosclerotic cardiovascular disease (ASCVD) and without (non-ASCVD); iPACE = Immuno-Proteomic Age of Cardiovascular Etiology.

The overall Pearson correlation between iPACE and cAge for the 1KIP cohort was 0.84 (p<0.001). Individual patient iPACE trajectories are shown in Figure 2B and calculated by the slope of serial measurements. The mean square errors (MSEs) for each year after enrollment, shown in Figure 2C, are fairly consistent (range: 4-7 years), suggesting model robustness. This positive but in-exact correlation between cAge and iPACE was predicted to capture the association of age-related immunoproteomic changes while still maintaining the variability of aging phenotypes that may be more or less resilient to CVDs and all-cause mortality. To this degree, we observed significantly higher iPACE residuals among patients who reported an ASCVD event throughout the duration of the study (n=380, Wilcoxon p<0.001, **Figure 2d**).

### iPACE is Correlated with Heart Failure in the INSPIRE cohort

To evaluate the robustness of the constructed age metric in a naive dataset, we calculated iPACE and iPACE residual for the INSPIRE participants (test dataset), and iPACE was positively correlated with cAge (R=0.38, p<0.001; MSE=7.6 years) (Figure 3A). Participants with HFpEF had significantly higher median iPACE residual than the case-matched control group (1.17 vs. -1.67, p<0.001, one-sided Wilcoxon test) (Figure 3B). Participants with HFpEF were more likely to have a history of renal failure compared to non-HFpEF participants (23.8 vs. 1.1%, p<0.001, Fisher’s exact test) and had higher median brain natriuretic peptide (BNP) (3.01 vs. 2.80, p<0.001, Wilcoxon rank-sum) and pro-BNP (4.94 vs. 4.38, p<0.001, Wilcoxon rank-sum) on baseline evaluation. iPACE residual was significantly correlated with pro-BNP in HFpEF patients (R=0.31, p<0.01) but not non-HFpEF (R=0.19, p=0.08) patients. While iPACE was not significantly correlated with BNP in participants with (R=0.07, p=0.51) or without (R=0.06, p=0.60) HFpEF.

**Figure 3.**
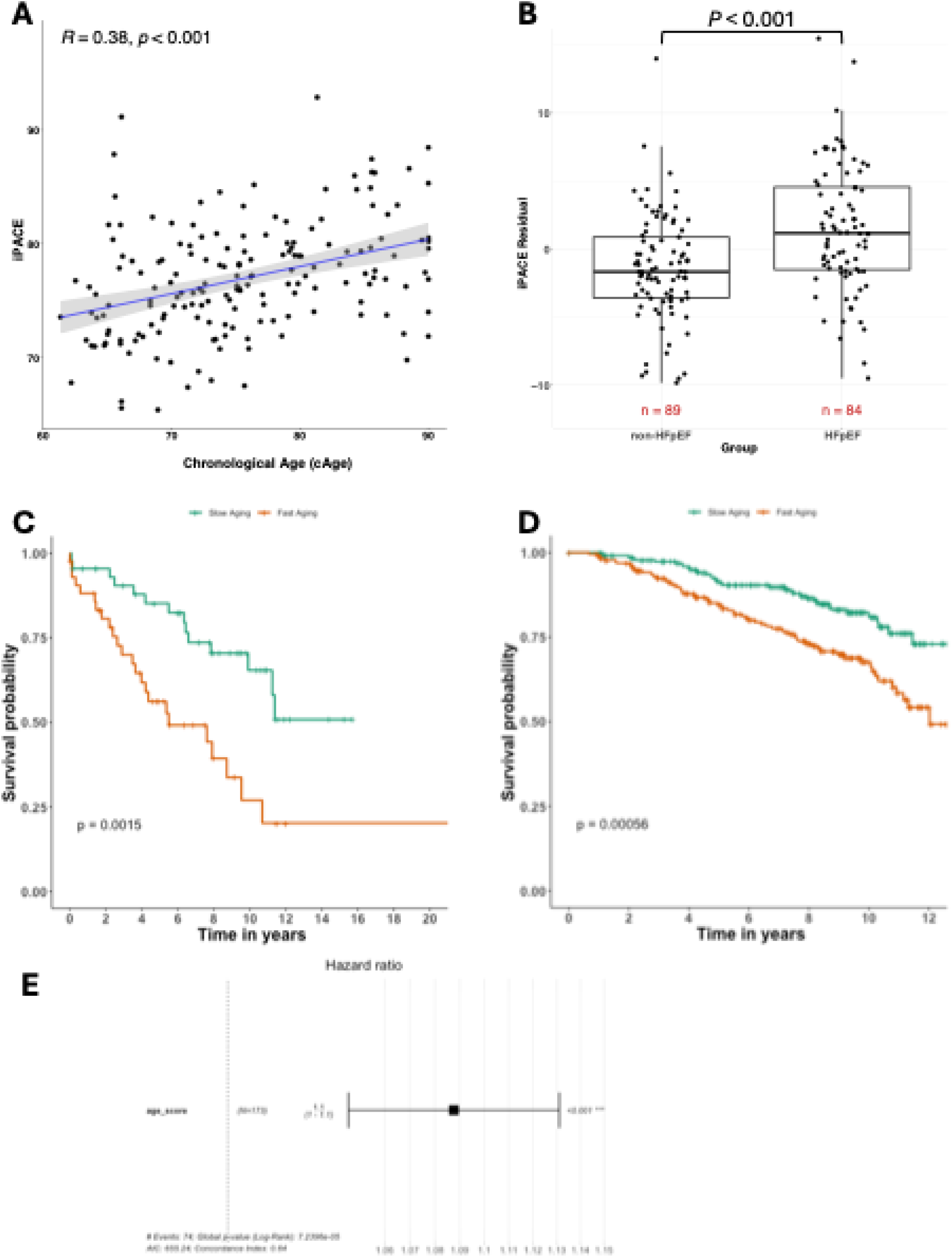
iPACE Model Validation. INtermountain Healthcare Biological Samples Collection Project and Investigational Registry (INSPIRE) [testing dataset] results: A) Pearson’s correlation coefficient of iPACE to cAge, B) iPACE residual by HFpEF status; C) Kaplan–Meier overall survival curves for Low vs. High iPACE participants in the INSPIRE cohort. D) Kaplan–Meier cardiovascular mortality curves for the Longevity (1KIP) cohort, color-coded to match INSPIRE. E) Cox proportional hazards regression model showing hazard ratios for all-cause and ASCVD outcomes by iPACE residual quartiles. HFpEF = heart failure with preserved ejection fraction; non-HFpEF = No heart failure with preserved ejection fraction; ASCVD = atherosclerotic cardiovascular disease, iPACE = Immuno-Proteomic Age of Cardiovascular Etiology.

### iPACE is associated with survival outcomes across independent cohorts

Among participants without HFpEF in the INSPIRE cohort, the Low iPACE group showed more favorable overall and ASCVD-free survival compared to the High iPACE group (Figure 3C). These results were further supported by survival analyses in the Longevity (1KIP) cohort, which demonstrated consistent trends for cardiovascular mortality (Figure 3D). In Cox proportional hazards regression, iPACE remained a significant predictor of all-cause and ASCVD-related mortality (Figure 3E).

## Discussion

In this study, we utilized serum proteomics of patients enrolled in large longitudinal studies to construct a. biological age metric. iPACE identifies thirty circulating immune biomarkers that may help unravel the age-related pathophysiology of HFpEF and susceptibility to ASCVD. The findings of this study build on previous research suggesting the development of HFpEF and ASCVD, to be linked to broad inflammatory perturbations [6,7] by further characterizing the aging component together referred to as “inflammaging” [8]. This is, to our knowledge, the first study to incorporate the use of proteomics to characterize biological age and its relationship to CVD.

Proteins identified as primary contributors to accelerated cardiovascular aging (iPACE coefficient>1) include PIGF, GDF-15, Adipsin, IL-12p40, and D-dimer. Placental growth factor (PlGF) is a member of the vascular endothelial growth factor (VEGF) family and is shown to be upregulated in atherosclerotic lesions in response to ischemia. Though still beginning to be uncovered, PlGF is thought to exhibit a protective role in ASCVD, promoting endothelial cell growth and survival in hypoxic conditions [9]. Growth/Differentiation Factor-15 (GDF15) is a member of the transforming growth factor-β cytokine superfamily and a marker of cell injury and inflammation [10]. Although the role of GDF15 in ASCVD and HFpEF remains unclear [11], a 2017 randomized control trial found circulating GDF15 levels to be independently associated with poor heart failure prognosis after controlling for conventional CVD risk factors and prognostic biomarkers [12].

Conversely, Leptin and ADAMTS were identified as primary contributors to delayed cardiovascular aging (iPACE coefficient<-1). Because leptin is adipocyte-derived, circulating concentrations are highly correlated with fat mass and may be confounded by obesity- and frailty-related conditions [13]. Therefore, it remains unclear if, how, and to what degree leptin independently influences age-related CVDs. Lastly, ADAMTS (a disintegrin and metalloproteinase with thrombospondin motifs) is a family of multidomain protease enzymes that regulate the remodeling of the extracellular matrix and are critical for versican (VCAN) degradation in the heart. In a recent study, impaired ADAMTS function contributed to excessive VCAN deposition and impairment of cardiac function, making the ADAMTS/VCAN axis an emerging target for drug development [14].

We acknowledge several limitations of this study. First, the analysis was performed in a retrospective fashion and thus was inherently limited in its ability to make conclusions about causality. Second, although we consider the multi-institutional nature of this study a strength by allowing us to construct two robust and longitudinal databases of distinct geographical origin, the collaboration was limited to large healthcare networks in Northern California, Southern Nevada, and Northern Utah, which may not be generalizable to all patient populations. Also, related to the longitudinal, multi-institutional nature of our study, general variability in practice patterns between and within institutions during the study period was expected and addressed by analyzing the impact of iPACE on CVD morbidity and mortality in the first year of the study. For these reasons, future clinical and experimental studies will be needed to externally validate the prognostic role of the cumulative iPACE signature and determine the pathophysiological role of individual iPACE proteins, respectively.

In conclusion, iPACE is an immunoproteomic signature of aging and validated as a predictor of age-related CVD (HFpEF and ASCVD) in a distinct cohort than the one from which it was derived. Accelerated aging was found to be associated with HFpEF and predictive of shortened all-cause mortality and ASCVD-free survival among patients without HFpEF. Nonetheless, future studies are required to verify the predictive power of iPACE and investigate whether therapies that modify iPACE translate to meaningful changes in CVD-related outcomes.

## METHOD

### iPACE Calculation and Correlation Analysis

To identify predictors of chronological age (cAge), we applied elastic net regression to circulating biomarker data from participants in the 1KIP cohort, which served as the training dataset. The elastic net model was implemented using the glmnet package in R. Prior to model training, all protein measurements were log-transformed and z-score standardized. Data collected during the initial year of enrollment were used to train the model, while data from subsequent years were reserved to evaluate the model’s robustness. To optimize the model parameters, we employed leave-one-out cross-validation across a 100 x 100 grid to determine the best values for the penalization parameter (λ) and the elastic net mixing parameter (α). The resulting iPACE values were derived as the fitted outputs from this optimized model. The optimized model was then used to determine iPACE for both 1KIP participants in the years following enrollment and for individuals in the independent INSPIRE testing cohort. Correlation analyses between iPACE and cAge, as well as other relevant phenotypic variables, were carried out using standard statistical methods in R.

### Survival Analysis for INSPIRE Participants

To investigate the prognostic utility of iPACE, we conducted survival analyses in the INSPIRE cohort, an independent validation population. Participants were stratified into three groups based on the residual difference between their iPACE and cAge, representing estimated biological age acceleration or deceleration. The groups were defined as “Low iPACE” (biologically younger than chronological age), “Normal iPACE” (minimal age discrepancy), and “High iPACE” (biologically older than chronological age).

We employed Kaplan-Meier survival analysis to estimate survival functions across these strata and applied Cox proportional hazards regression to quantify associations between iPACE-based categorization and health outcomes, including ASCVD and all-cause mortality. All survival analyses were conducted using the survival package in R, including parallel analyses for the Longevity (1KIP) and INSPIRE cohorts.

## DATA AVAILABILITY

This study analyzed existing data from the Stanford 1000 Immunomes Project (1KIP) and INSPIRE cohorts, collected independently of this study.

## AUTHOR CONTRIBUTIONS

All authors contributed to the writing of this manuscript.

## ACKNOWLEDGEMENTS

National Institutes of Health grant R01DK128435 (D.A.W.), and Canadian Institutes of Health grant PJT-186165 (D.A.W.), the Buck Institute for Research on Aging Bioinformatics and Data Science Core (F.W., J.B., M.F., D.F.), National Institute on Aging grants: P01AI153559 (F.W., D.F.), 5T32AG000266-23 (J.B.).

